# GIBBERELLIN SIGNALING THROUGH RGA SUPPRESSES GCN5 EFFECT ON STAMEN ELONGATION OF ARABIDOPSIS FLOWERS

**DOI:** 10.1101/2024.04.30.591935

**Authors:** Christina Balouri, Stylianos Poulios, Dimitra Tsompani, Zoe Spyropoulou, Maria-Christina Ketikoglou, Athanasios Kaldis, John H Doonan, Konstantinos E Vlachonasios

## Abstract

Histone acetyltransferases (HAT) modify the amino-terminal tails of the core histone proteins via acetylation, regulating chromatin structure and transcription. The GENERAL CONTROL NON-DEREPRESSIBLE 5 (GCN5) is a HAT that specifically acetylates H3K14 residues. GCN5 has been associated with cell division and differentiation, meristem function, root, stem, foliar and floral development, and plant environmental response. The flowers of *gcn5–6* plants display reduced length of stamen and exhibit male sterility relative to the wild-type plants. We show these effects may arise from gibberellin (GA) signaling defects. The signaling pathway of bioactive GAs depends on the proteolysis of their repressors, DELLA proteins. The DELLA protein, REPRESSOR OF GA (RGA), represses plant growth, inflorescence, flower and seed development. Our molecular data indicate that GCN5 is required for activation and H3K14 acetylation of genes involved in the late stages of GA biosynthesis and catabolism. We studied the genetic interaction of RGA and GCN5; RGA can partially suppress GCN5 action. The reduced elongation of the stamen filament of *gcn5–6* mutants is reversed in the *rga–t2;gcn5–6* double mutants. This mechanism involved suppressing the GCN5 effect on the expression and histone acetylation in *GAI*-locus by RGA. Interestingly, RGA and RGL2 do not suppress ADA2b function, suggesting that ADA2b acts downstream in GA signaling and is distinct from GCN5 activity. In conclusion, we propose that the action of GCN5 on stamen elongation is mediated partially by RGA and GA signaling.

## Introduction

For activation of gene expression during development, transcription factors must overcome repressive chromatin structure, which is accomplished with the help of multiprotein complexes (Hegde and Bernstein 2006; Li et al. 2007). One class of complexes modify the nucleosomal histones through acetylation, phosphorylation, methylation and other modifications (Bannister and Kouzarides 2011). Acetylation of specific lysine residues in histone N-terminal tails is catalyzed by Histone AcetylTransferases (HAT), which are involved in transcriptional regulation and other nuclear processes. HATs are part of large multiprotein complexes, like the SAGA complex, in which their activity is enhanced, their substrate specificity is modified, and the whole complex is recruited to target sequences on the genome with the help of other components involved in protein-protein interactions (Grasser et al. 2021). HATs and HDACs (histone deacetylases) can target promoters to activate or suppress gene expression, respectively.

GENERAL CONTROL NON-DEREPRESSIBLE 5 (GCN5, also known as HAG1) is a histone acetyltransferase (Stockinger et al. 2001; Pandey et al. 2002) involved in many developmental processes and responses to environmental stimuli (Grasser et al. 2021; Vlachonasios et al. 2021). The *gcn5* mutants exhibit pleiotropic phenotypes, including dwarfism, loss of apical dominance, upward curled and serrated leaves, abnormal inflorescence meristem, abnormal flower development, and shorter roots (Bertrand et al. 2003; Vlachonasios et al. 2003; Cohen et al. 2009; Poulios and Vlachonasios 2016; Poulios et al. 2022; Tsilimigka et al. 2022). GCN5 is the catalytic subunit of the Arabidopsis SAGA (Spt-Ada-GCN5-acetyltransferase) complex (Servet et al. 2010; Grasser et al. 2021; Vlachonasios et al. 2021; Wu et al. 2021) that acetylates H3 and H2A in nucleosomes and has been shown to interact with the transcriptional adaptors ADA2a and ADA2b (Stockinger et al. 2001; Mao et al. 2006; Poulios et al. 2021). The *ada2b* mutants present a phenotype highly similar but not identical to *gcn5* mutants (Sieberer et al. 2003; Vlachonasios et al. 2003; Hark et al. 2009). The *ada2b* mutants are characterised by dwarfism, small dark green curled leaves and infertility (Vlachonasios et al. 2003). GCN5 and ADA2b affect gynoecium development by modulating auxin and cytokinin signaling (Poulios and Vlachonasios 2018). Furthermore, loss of function GCN5 affects stamen elongation, especially in flowers formed early in development (Cohen et al. 2009).

Plant hormones, specifically gibberellins, modulate stamen development and function (Plackett et al. 2011). Gibberellins (GAs) biosynthetic mutants revealed that GAs stimulate stamen filament elongation through increased cell elongation and promote anther dehiscence (Marciniak and Przedniczek 2019). In Arabidopsis, GA is perceived by three GA receptors: Gibberellin Insensitive Dwarf 1s (GID1a, GID1b and GID1c). The triple GA insensitive mutant produces a more severe stamen phenotype than the mutant in GA biosynthesis, *ga1-3* (Griffiths et al. 2006). The binding of GA to these receptors promotes the interaction with DELLA proteins, which are GRAS domain proteins and major repressors of GA signaling (Sun 2010). In Arabidopsis, five DELLA proteins have been identified: GA Insensitive (GAI), Repressor of GA1-3 (RGA) and RGA Like (RGL1, RGL2 and RGL3). The binding of DELLA proteins to the GA-GID1 complex results in polyubiquitination and triggers their degradation by 26S proteasome (Hirano et al. 2008). Several transcription factors have been identified downstream of DELLA proteins, including members of PIFs and MYB families (Cao et al. 2006; de Lucas et al. 2008). Specifically, MYB21 and MYB24 control stamen filament growth by acting downstream of DELLAs (Peng 2009).

In this study, we explored the role of GCN5 in gibberellin responses and demonstrated using genetic and molecular approaches that RGA partially suppress the GCN5 effect on stamen elongation by affecting the H3 acetylation on genes involved in gibberellin biosynthesis, catabolism and signaling.

## Results

### ADA2a, ADA2b and GCN5 are required for the hypocotyl response to exogenous GA

Hypocotyl elongation is affected in SAGA-related mutations (Vlachonasios et al. 2003) and is a GA-sensitive process. Therefore, we hypothesised that mutant hypocotyls might display altered responses to exogenous GA. The effect of gibberellins on hypocotyl elongation of *ada2a*, *ada2b* and *gcn5* mutant seedlings was measured for five consecutive days after applying 10 µM GA_3_. Initially, applying gibberellins increased the hypocotyl growth in both Ws-2 and Col-0 wild-type seedlings (Fig. 1A and D). The response was more significant in Col-0 seedlings than in Ws-2. The hypocotyl growth in *gcn5-1* and *gcn5-6* mutants was slower than in wild-type plants, and the response to gibberellins was minor and measurable after four days of exposure (Fig. 1B and E). The sensitivity of both *gcn5* mutants to GA_3_ was reduced (Fig. 1H and I), suggesting that GCN5 is required for GA-induced hypocotyl elongation. The response of two *ada2b* mutant alleles, *ada2b-1* and *prz1-1,* to gibberellins was also lower than that of the wild-type plants but with a shorter delay of two days (Fig. 1C and F). The sensitivity to GA_3_ of both *ada2b* mutants was also reduced (Fig. 1H and I), implying that ADA2b is also essential for GA-induced hypocotyl growth. In contrast, *ada2a-3* mutants displayed an increased hypocotyl elongation upon GA treatment, albeit lower than wild-type plants (Fig. 1G). The sensitivity of *ada2a-3* was higher than *gcn5* and *ada2b* mutants but lower than the wild-type plants (Fig. 1I), suggesting that ADA2a has a minor role in GA-induced hypocotyl growth.

**Figure 1.**
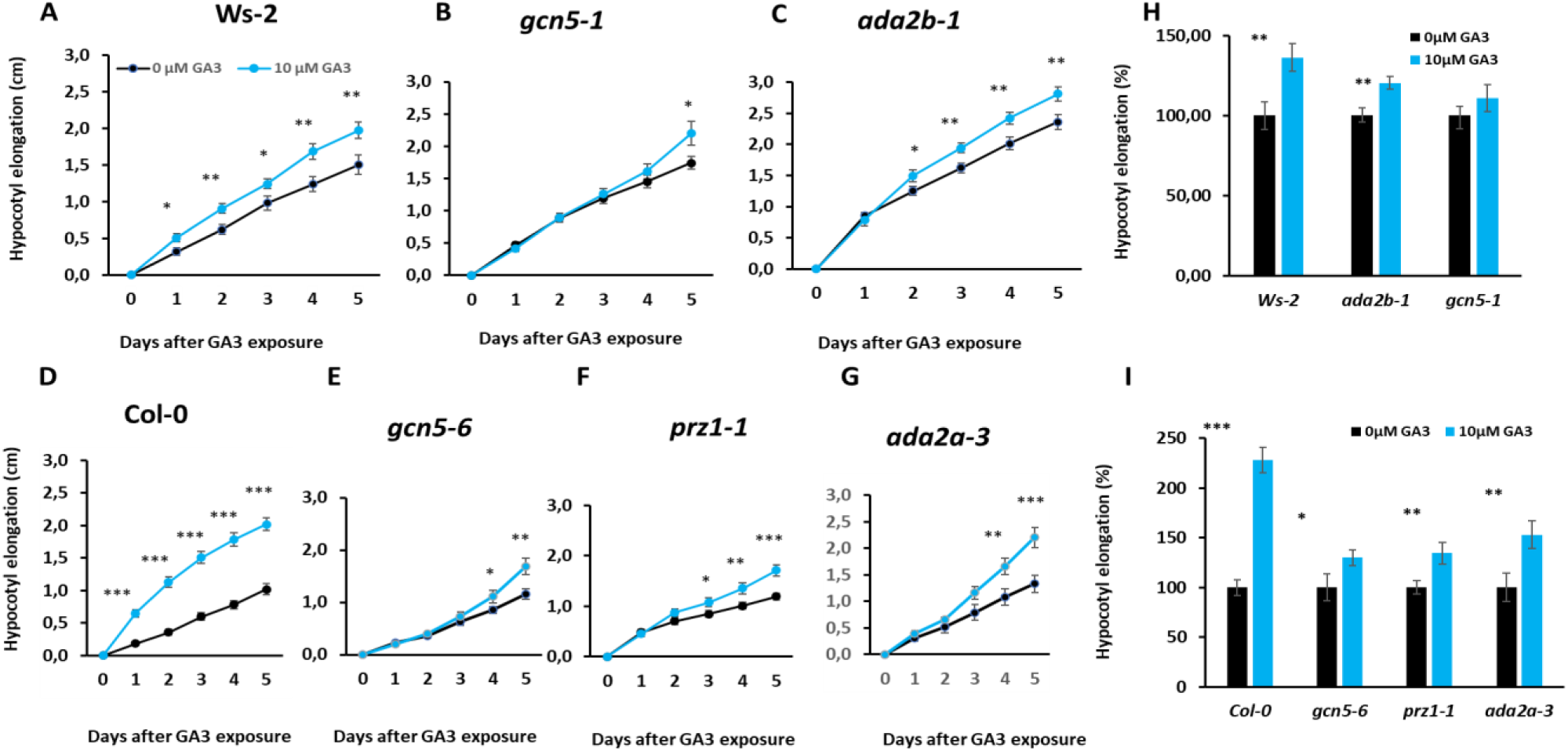
The effect of GA_3_ on hypocotyl elongation of Ws-2 (A), *gcn5-1* (B), *ada2b-1* (C), Col-0 (D), *gcn5-6* (E), *prz1-1* (F) and *ada2a-3* (G) seedlings. Sensitivity of Ws-2, *gcn5-1* and *ada2b-1* (H), and Col-0, *gcn5-6*, *prz1-1* and *ada2a-3* (G) mutant seedlings after four days of exposure to 10μΜ GA_3._ Asterisks *, ** and *** indicate statistically significant differences from 0μΜ GA_3_ using Student’s t-test for P< 0.05, P < 0.01 and P < 0.001, respectively (n=90 plants).

### The role of ADA2a, ADA2b and GCN5 in the root elongation of seedlings after exposure to gibberellins

Root growth is affected by GCN5 and ADA2b (Vlachonasios et al. 2003; Kornet and Scheres 2008). Root elongation depends on the action of gibberellins (Barker et al. 2021). The effect of gibberellins on primary root elongation of *ada2a*, *ada2b* and *gcn5* mutant seedlings was measured for 4 or 5 consecutive days after applying 10 µM GA_3_. The exposure to gibberellins slightly decreased the root growth in both Ws-2 and Col-0 wild-type seedlings (Supplemental Fig. S1A and E). Although the roots of *ada2b-1* and *prz1-1* are short, the response to gibberellins was similar to that of the wild type (Supplemental Fig. S1B and F). In contrast, *gcn5-1* and *gcn5-6* mutants were more responsive to gibberellin. Despite the short root, their exposure to gibberellins reduces the root elongation of *gcn5* mutants (Supplemental Fig. S1C and G). The sensitivity of root elongation to gibberellins was higher in the *gcn5* mutants suggesting that GCN5 is a negative regulator of GA-induced root growth retardation (Supplemental Fig. S1I). Finally, the root growth of *ada2a-3* seedlings decreased upon exposure to GA3, similarly to *gcn5* mutants indicating that ADA2a is also implicated as a negative regulator of GA-induced root growth retardation (Supplemental Fig. S1H and I).

### ADA2b and GCN5 affect flower morphology by modulating gibberellin biosynthesis

Arabidopsis mutants in *GCN5* and *ADA2b* are characterised by abnormal flower development (Vlachonasios et al. 2003). Both mutant plants displayed short stamens, especially in the early-formed flowers (Fig. 2A). The stamen elongation is restored only in the late-forming *gcn5* flowers (Vlachonasios et al. 2003; Cohen et al. 2009). Defects on gibberellin biosynthesis or signaling could raise this effect (Griffiths et al. 2006). We, therefore, monitored the expression of the *GA3ox1*, the last enzyme in the GA biosynthesis, in the *ada2b-1* and *gcn5-1* early-formed flowers. Indeed, the expression of *GA3ox1* was dramatically reduced in both mutants (Fig. 2B), suggesting that GCN5 and ADA2b act as positive regulators of GA biosynthesis gene expression in early-formed flowers in Arabidopsis. As a result, we hypothesised that GCN5 and ADA2b regulate GA biosynthesis in flowers through GA signaling components, including DELLA proteins. The DELLA proteins, RGA, RGL1 and RGL2, are known to be required for flower development (Daviere and Achard 2013) and to be involved in GA biosynthesis (Zentella et al. 2007). Therefore, a genetic approach was carried out by crossing *gcn5-6 (hag1-6)* and *ada2b-1* mutants with *rga* mutants to test the proposed regulation in flowers and hypocotyl elongation.

**Figure 2.**
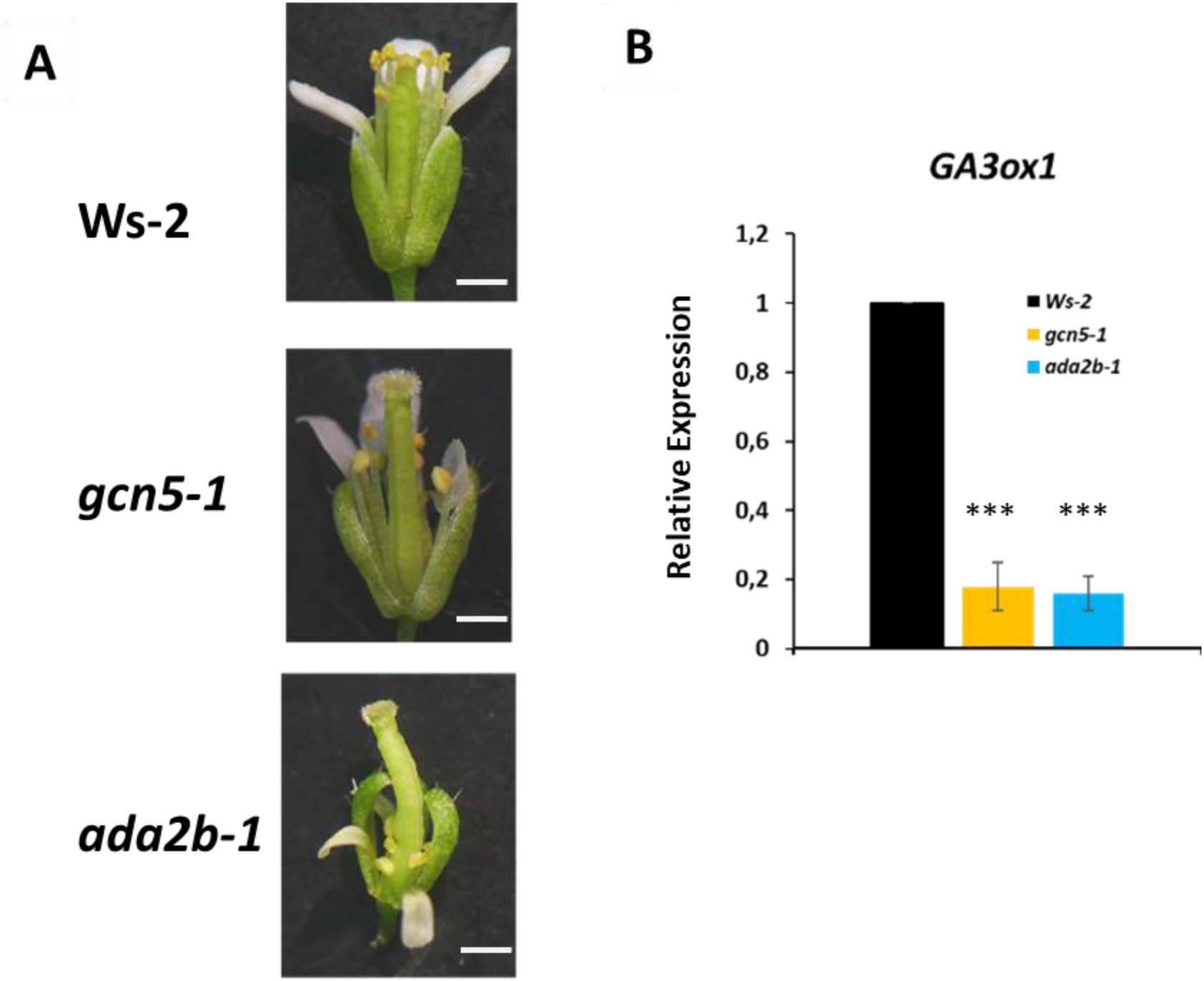
GCN5 and ADA2b affect flower development. Α) Early-flower formed in *gcn5-1*, *ada2b-1* and wild-type Ws-2, Bar equals 0.5 cm. Β) *GA3ox1* expression in flowers of *gcn5-1*, *ada2b-1* and wild-type. Asterisks *** indicate statistically significant differences from Ws-2 for P < 0.001, respectively.

### Mutation in *RGA* partially suppresses *gcn5* phenotypes

#### Hypocotyl elongation is restored in *rga;gcn5* double mutants

In young seedlings, the *gcn5–6* mutant showed reduced hypocotyl elongation (Fig. 3A and B), whereas an elongated hypocotyl was observed in the *rga-t2* seedlings, compared to the wild type (Col-0). In contrast, in the *rga–t2;gcn5–6* double mutant, the hypocotyl phenotype resembled *rga–t2*, completely reversing the hypocotyl length of the *gcn5–6* mutant, suggesting that RGA could suppress GCN5 effect on hypocotyl elongation. The root growth in the *gcn5–6* mutant was significantly reduced compared to wild-type and *rga-t2* mutant plants (Fig. 3A and C). In the *rga–t2;gcn5–6* mutant, the root length was similar to *gcn5* root growth. Therefore, RGA represses the GCN5 action on hypocotyl elongation in the light, acting downstream of GCN5. In contrast, GCN5 promotes root elongation independently from RGA.

**Figure 3.**
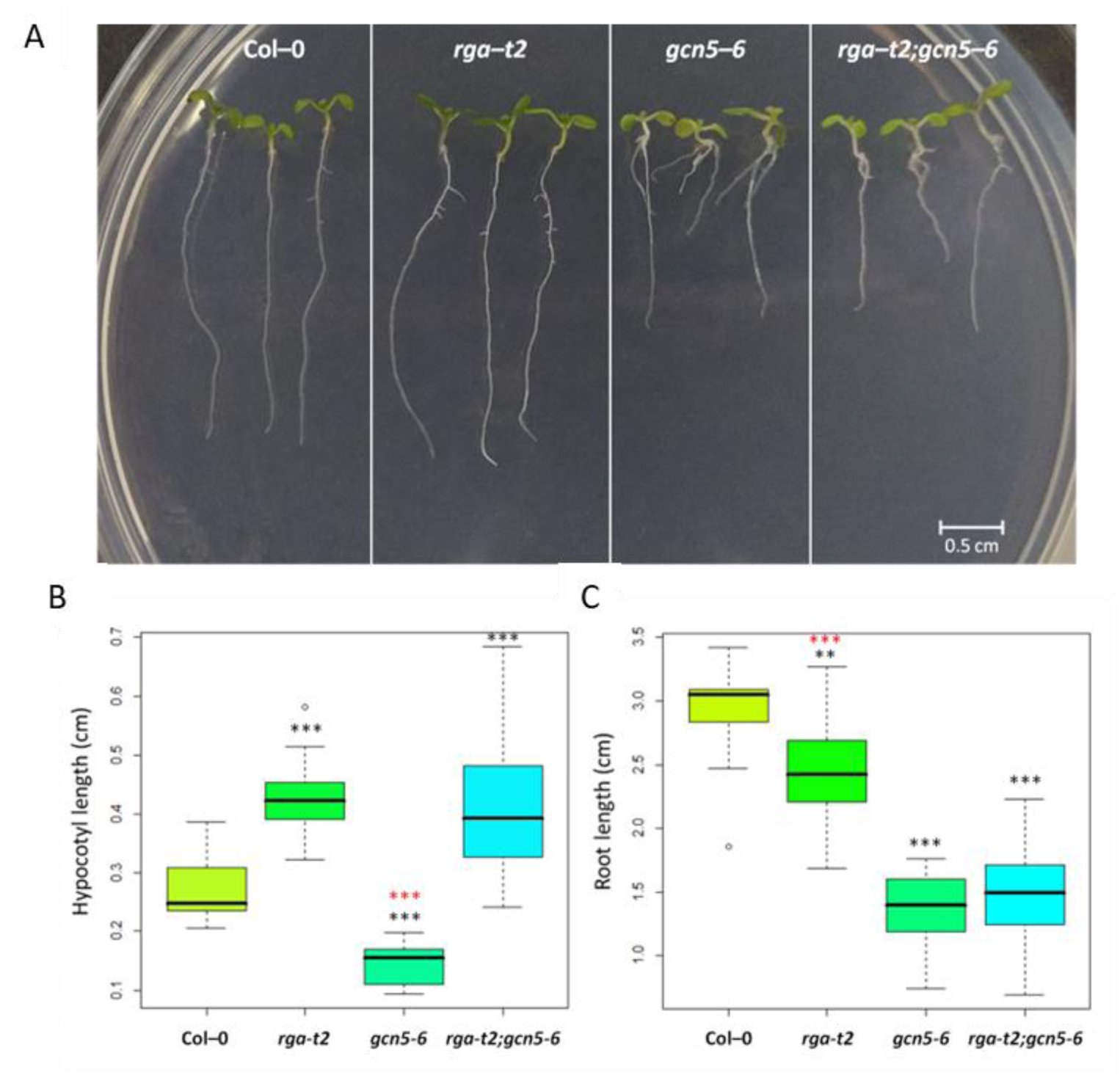
Characterisation of *rga-t2;gcn5-6* double mutants at seedlings stage. (A) Seedlings of Col–0, *rga–t2*, *gcn5–6* and *rga–t2; gcn5–6*, at day 7 of growth. The length of the hypocotyl (B) and root (C) of seven-day-old seedlings. Asterisks ** and *** indicate statistically significant differences from Col–0 for P < 0.01 and P < 0.001, respectively. Similarly, divergence from the *rga–t2;gcn5–6* double mutant is shown with red asterisks. Error bars represent the standard deviation.

#### Loss of RGA did not suppress gcn5 leaf phenotype

During the vegetative stage, the *gcn5–6* mutant exhibits a distinct phenotype with small serrated leaves. Interestingly, even after 20 days of growth, the rosette leaves of the *rga–t2;gcn5–6* double mutant plants remained serrated, indicating that RGA does not play a significant role in the GCN5 effect on leaf development (Fig. 4).

**Figure 4.**
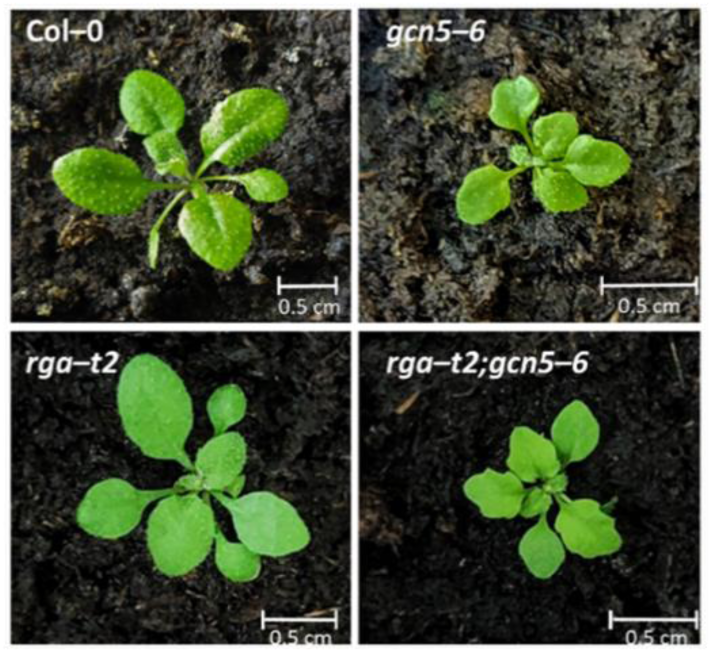
The effect of GCN5 and RGA on leaf development. Rosette development of twenty days old Col-0, *gcn5-6*, *rga-t2* and *rga-t2;gcn5-6* plants.

#### GCN5 is required for the inflorescence growth and is partially suppressed by loss of RGA

In later development, *gcn5–6* mutant plants are characterised by delayed flowering as previously described (Kim et al. 2015) and short inflorescence relative to wild-type Col–0 plants. In contrast, *rga–t2* plants flower earlier and show longer inflorescence than wild-type plants. In the *rga–t2;gcn5–6* double mutant plants, the inflorescence growth is partially restored, compared to *gcn5–6*, without reaching the growth rate of Col–0 (Fig. 5A). After the opening of the first flower, at 30–35 days of age for the wild type, *rga* and *rga–t2;gcn5–6* mutants and ∼50 days for the *gcn5–6* mutant, the number of lateral inflorescences and the length of internodes were measured. Two lateral inflorescences are identified in wild-type plants and the *gcn5–6* mutant. The *rga–t2* and *rga–t2;gcn5–6* mutants have 2 or 3 lateral inflorescences, suggesting that RGA regulates the number of lateral inflorescences in Arabidopsis (Supplemental Table S1). As shown in Fig. 5B, the length of the first internode, which refers to the basal part of the inflorescence starting from the rosette to the first axillary bud, is shorter in *gcn5-6* mutant than Col-0 and *rga–t2* mutant plants. In the *rga–t2;gcn5–6* double mutant, the length of the first internode is noticeably shorter than Col–0 and *rga–t2*, while it does not show a statistically significant difference from the *gcn5–6* mutant. Therefore, *rga-t2* can not suppress the *gcn5* defect for the first internode. The second internode, which concerns the part of the inflorescence between the first and second cauline leaf, is longer in the *rga–t2* compared to Col–0, while *gcn5–6* again shows a reduction in length. In the double mutant, however, the length of the second internode is greater than in *gcn5–6* mutant plants (Fig. 5C). These results suggest that GCN5 is required to elongate the internodes in the inflorescence growth properly and is partially suppressed by RGA. A third internode is found only in the *rga–t2* and *rga–t2;gcn5–6* mutants. The length of the third internode appears to be slightly elongated in the double mutant (Fig. 5D).

**Figure 5.**
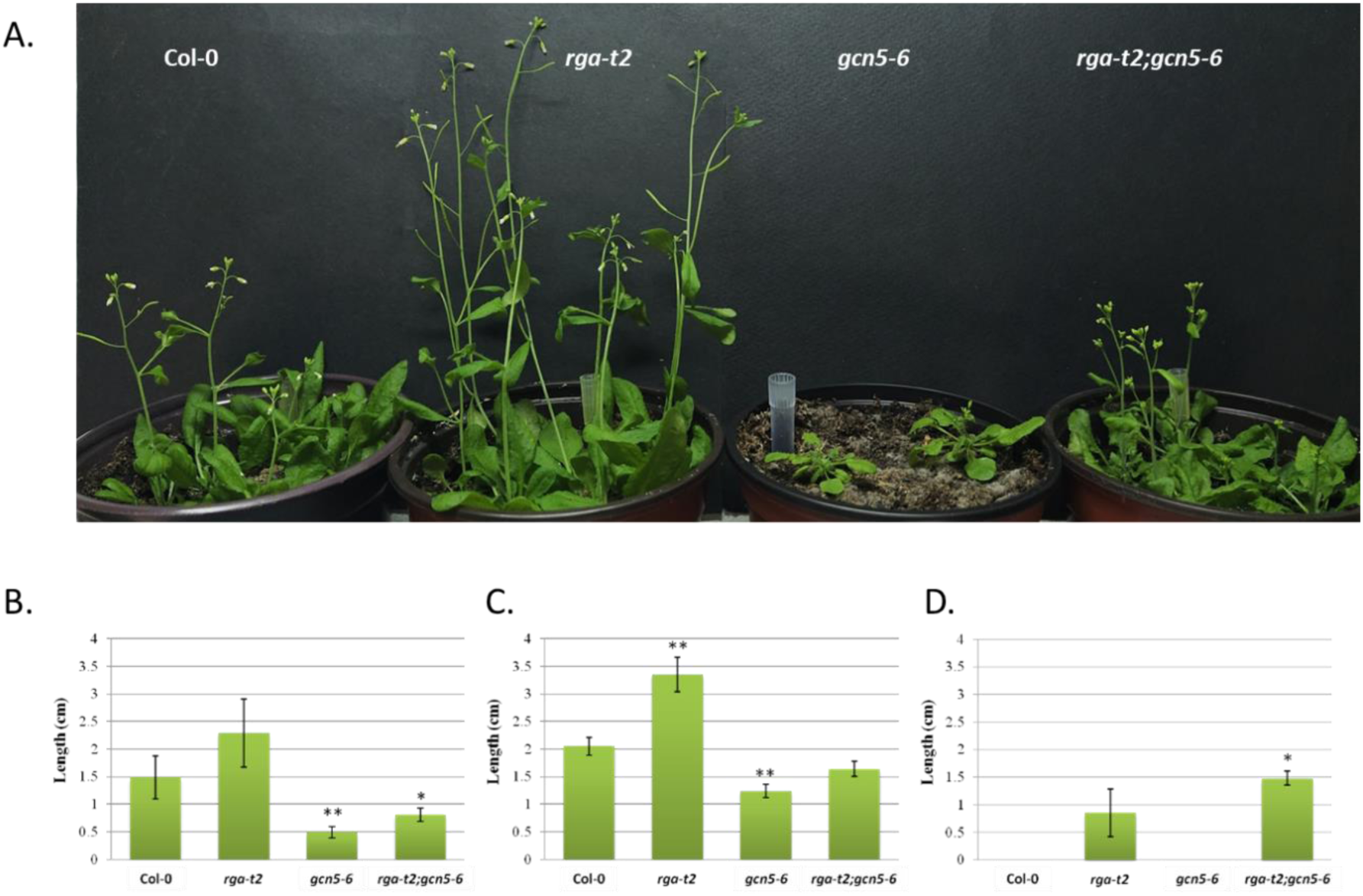
Col-0, *rga-t2*, *gcn5-6* and *rga-t2gcn5-6* mutant plants after 36 days of growth (A). The effect of RGA and GCN5 on first internode length (B), second internode length (C) and third internode length (D). Asterisks * and ** in B and C indicate statistically significant differences from Col–0 for P < 0.05 and P < 0.01, respectively and in D indicates a significant difference from *rga-t2* for P < 0.05. Error bars represent the standard deviation (n=16).

After two months, Col–0, *rga–t2* and double mutant plants completed their life cycle while the *gcn5–6* mutant continued growing. The final height of *rga–t2* plants is significantly higher than Col–0. In contrast, the *gcn5–6* mutant plants are still flowering, extending their lifespan. The *rga–t2;gcn5–6* double mutant shows a slightly longer life cycle than wild-type plants but shorter than *gcn5–6.* Its final main inflorescence length reaches that of the wild type, completely reversing the developmental problem observed in the *gcn5–6* mutant plants (Supplemental Fig. S2 and Fig. S3A), suggesting that RGA is required for the GCN5 action in the latest stages of plant development. At the third month of growth, *gcn5–6* plants do not grow in height; however, they become very bushy (Supplemental Fig. S3B) due to numerous secondary inflorescences (Supplemental Table S2). This phenotype is reversed in the double mutant, indicating that RGA is required for the GCN5 regulation on secondary inflorescence formation.

#### RGA is required for GCN5 function on stamen elongation in early flowers

During inflorescence development, the early flowers of the *gcn5–6* mutants have short petals that do not exceed the length of the sepals (Fig. 6E and F). The number of stamens remains the same (6) as in the wild type (Supplemental Table S3). However, the length of the stamen filament is significantly shorter (Fig. 6I), associated with reduced fertility, suggesting that GCN5 is a positive regulator of stamen filament growth. In the *rga-t2* mutant, the number of stamens varies between 5 and 6 (Supplemental Table S3) and the length of the stamen filament is increased compared to wild-type flowers (Fig. 6A, B, C, D, I). In the double mutant, the petal length phenotype of *gcn5–6* is wholly reversed (Fig. 6G and H). The number of stamens (Supplemental Table S3) and filament length (Fig. 6I) displayed significant variation, but most flowers had more stamens than Col–0 andfilament length was fully restored, to wild-type. These results indicate that the removal of RGA fully compensates for the loss of GCN5 on stamen elongation.

**Figure 6.**
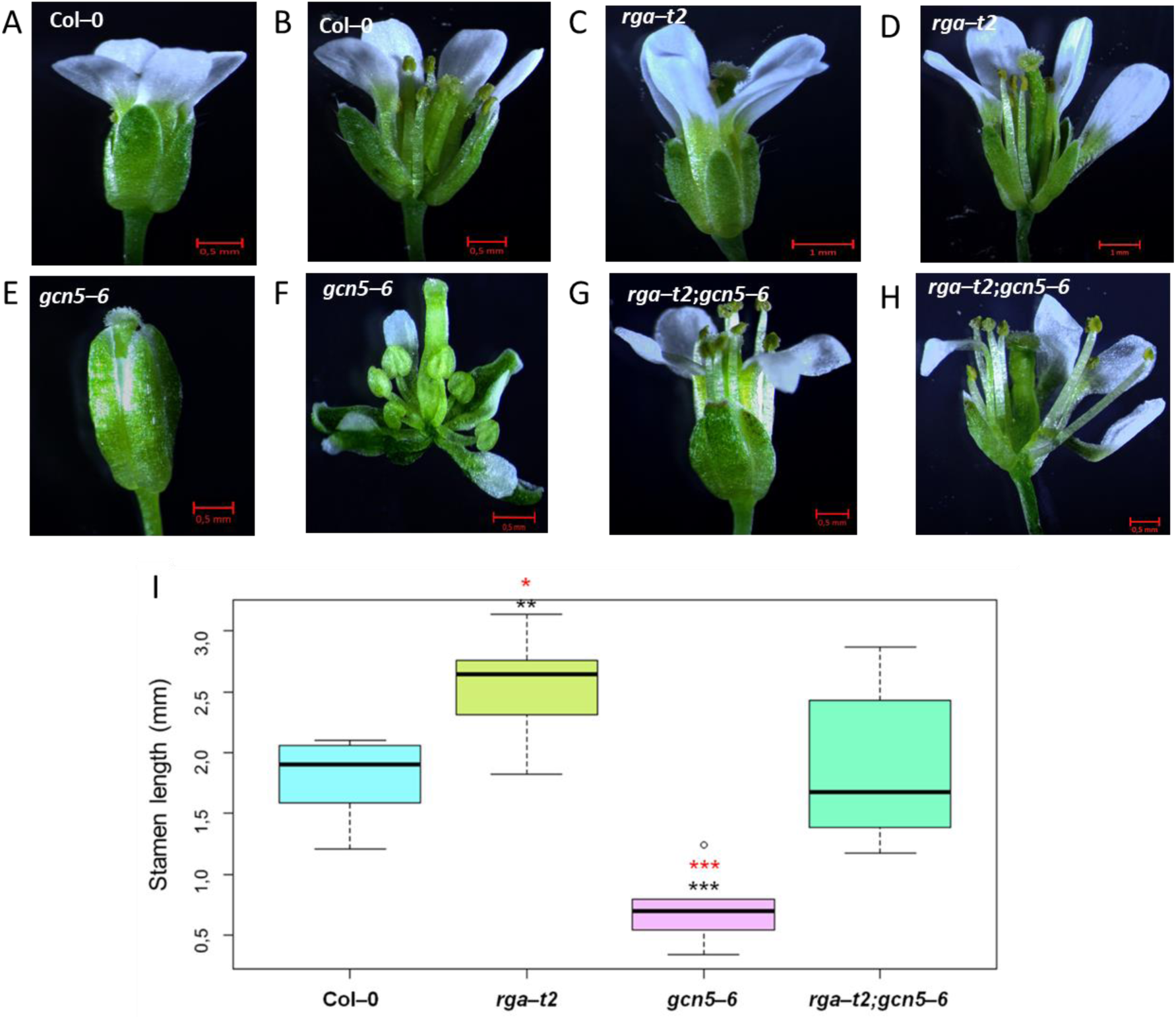
The effect of *GCN5* and *RGA* on first-formed flowers. Closed and open flower Col–0 (A and B), *rga–t2* (C and D), *gcn5–6* (E and F) and *rga–t2;gcn5–6* (G and H). Stamen filament length (I). Asterisks *, ** and *** indicate the statistically significant difference in filament length from Col–0 for P < 0.05, P < 0.01 and P < 0.001, respectively. Similarly, red asterisks show statistically significant differences from the stamen growth of the double mutant. Error bars represent the standard deviation.

The *gcn5–6* plants showed male sterility, which prevents the flower from self-fertilising. Effectively, no siliques and seeds are produced. In contrast, silique development and seed production are achieved in the *rga–t2;gcn5–6* double mutant. This result suggests that the removal of RGA can compensate for the loss of GCN5, leading to successful silique growth and seed production. However, the double mutant siliques are significantly shorter than those of the wild-type and *rga* mutants, suggesting a positive role of GCN5 on silique growth or pollen fertility (Supplemental Fig. S4).

### The effect of *GCN5* and *RGA* on gene expression in hypocotyls and flowers

Then, to further explore the effect of GCN5 and RGA in hypocotyl growth, we monitored the expression of genes involved in biosynthesis, catabolism and signaling of gibberellins in the hypocotyl and cotyledons. *GA3ox1*, the last gene in the GA biosynthesis pathway, which is highly expressed in cotyledons and the apical meristem (Mitchum et al., 2006), showed increased expression in *gcn5–6*, while in the *rga–t2;gcn5-6* double mutant, the expression was partially reversed (Supplemental Fig. S5A). The GA2ox family of proteins participates in the catabolism of gibberellins. It is known that the *GA2ox4* gene is expressed in the shoot apical meristem, the *GA2ox6* in the stem and vessels, and the *GA2ox8* is expressed in the leaf stomatal cells (Li et al. 2019). *GA2ox4* expression was increased in all mutants tested, with *gcn5–6* showing a four-fold increase from the wild type. In *rga–t2;gcn5– 6*, the *GA2ox4* expression was reversed close to the levels of the *rga–t2* mutant (Supplemental Fig. S5B). Expression of the *GA2ox6* gene did not change significantly in the *rga–t2* and *gcn5–6* mutants from the wild type, while in the double mutant, an increase was observed, suggesting a synergistic action of the RGA and GCN5 (Supplemental Fig. S5C). Finally, the *GA2ox8* gene displayed an increased expression in *gcn5–6* and the double mutant *rga– t2;gcn5–6* (Supplemental Fig. S5D). The GA receptor, *GID1b*, showed a slightly increased expression in the single mutants and a three-fold upregulation in *rga–t2;gcn5–6* (Supplemental Fig. S5F), while no significant difference was observed in the expression of *GID1a* between the genotypes (Supplemental Fig. S5E). *GAI* expression in rosette leaves was slightly decreased in *gcn5-1* and *ada2b-1* mutants, while there is no detectable change in *RGA* expression (Vlachonasios et al. 2003; Trachtman et al. 2019) In hypocotyls, the expression of *GAI* was slightly increased in the single and double mutants compared to wild-type (Supplemental Fig. S5G). *RGL2* showed reduced expression in the *gcn5–6* mutant, which was reversed in the double mutant and upregulated compared to the wild type and the *rga–t2* mutant (Supplemental Fig. S5H).

Using Real-time RT-PCR, we studied the gene expression in gibberellin biosynthesis, catabolism, and signaling in the early flowers. As shown in Fig. 7A, the *GA20ox2* is expressed at a lower level in the flowers of *gcn5–6* mutants in comparison with Col–0 and *rga–t2*, while in *rga–t2;gcn5–6,* a partial restoration of expression levels is found, but not significant. The *GA3ox1* gene is expressed in the base of the young flower, apex and sepal vessels and mainly in the filament of the stamens of the mature flower (Mitchum et al. 2006; Hu et al. 2008). As shown in Fig. 7B, a decreased expression in *gcn5–6* flowers was observed, whereas it has increased expression in *rga–t2* flowers compared to the wild type. In the double mutant, the expression of *GA3ox1* was higher than *gcn5-6*, close to wild-type levels. Therefore, RGA appears to reverse the expression levels of the *GA3ox1* in the *gcn5* background. Regarding the genes expressing GA catabolism enzymes, *GA2ox7* and *GA2ox8* are expressed in the whole flower (Li et al. 2019). The *GA2ox7* gene is not expressed in *gcn5–6* early-formed flowers (Fig. 7C). In the double mutant, *GA2ox7* expression is detectable at a lower level than wild-type or *rga* mutants. Similarly, *GA2ox8* expression is deficient in *gcn5–6* flowers compared to wild-type or *rga* mutants, while it is increased slightly in *the rga–t2; gcn5–6* double mutant (Fig. 7D). The expression of the GA receptor, *GID1b,* increased in the *gcn5-6* mutants compared to wild-type plants. In the *rga* mutant early flowers, *GID1b* was decreased to low levels. A decreased expression of *GID1b* in the double mutant was also observed (Fig. 7E). Therefore, RGA reverses the effect of GCN5 on GID1b expression levels. Reduced gene expression of the DELLA protein GAI is observed in the early-formed flowers of the *gcn5–6* mutant. The *GAI* expression in the flowers of *rga– t2;gcn5–6* mutants does not show a statistically significant deviation from the wild type (Fig. 7F). Therefore, the positive regulation of *GAI* expression by GCN5 in primary flowers appears to be reversed by the action of the DELLA protein RGA.

**Figure 7.**
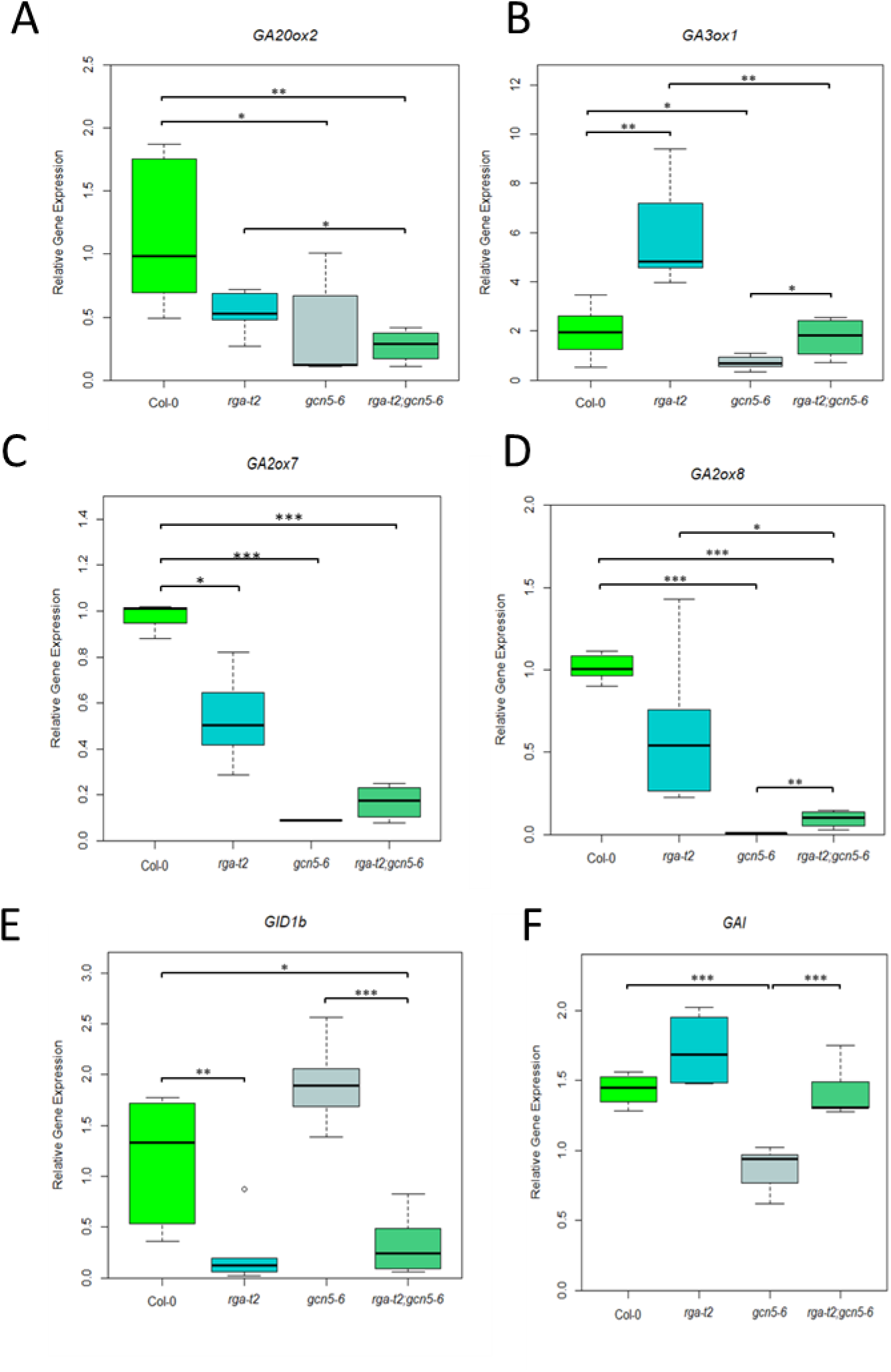
*Expression of gibberellin biosynthesis genes GA20ox2* (A) and *GA3ox1* (B), gibberellin catabolism genes *GA2ox7* (C) and *GA2ox8 (D),* GA-receptor *GID1b (E),* and DELLA gene *GAI (F)* in early flowers of *Arabidopsis thaliana* inflorescence. Error bars show the standard deviation. Asterisks show statistical significance compared to *Col–0* and *rga–t2;gcn5–6* double mutant, using Student’s t-test: *, P < 0.05, **, P < 0.01, and ***, P < 0.001.

#### GCN5 and RGA alter H3K14 acetylation in the promoter of GA-related genes

To examine whether the observed changes in gibberellin-related gene expression in the *gcn5-6* and the *rga-t2;gcn5-6* double mutant resulted from changes in the acetylation status of its locus, we performed ChIP analysis using antibodies for total histone H3 and acetylated lysine 14 in histone H3 (H3K14); H3K14 is known as the GCN5 target for acetylation (Benhamed *et al*. 2006, Earley *et al*. 2007). Our results showed that in the promoter region *GA20ox2* and *GA3ox1* loci, total histone H3 acetylation is reduced significantly in *gcn5-6* compared to wild-type plants (Fig. 8A and B). The double mutant’s H3K14 acetylation levels were also low, suggesting that H3K14 acetylation levels correlate with gene expression profile. Thus, the results suggest that GCN5 is required for the H3K14 acetylation of late GA biosynthetic genes, consistent with the altered expression levels. Then, we explored the effect of GCN5 and RGA on the H3 acetylation in two genes involved in GA-catabolism, *GA2ox7* and *GA2ox8*. The H3K14 acetylation level was almost undetectable in the *gcn5-6* plants compared to wild-type and *rga-t2* plants (Fig. 8C and D). In the double mutants, the level of H3K14 acetylation was restored to wild-type levels. These results suggest that H3K14 acetylation in the GA-catabolism genes is affected through the RGA pathway. A similar scenario is observed in the 5-UTR region of DELLA protein *GAI*; the level of H3K14 acetylation is low in the *gcn5-6* inflorescences compared to wild-type and *rga-t2* (Fig. 8E). In the double mutant, H3K14 acetylation returns to wild-type levels, again correlated with the gene expression profile. This result suggests that the loss of RGA action restores the GCN5 requirement for both expression and histone acetylation of GAI.

**Figure 8.**
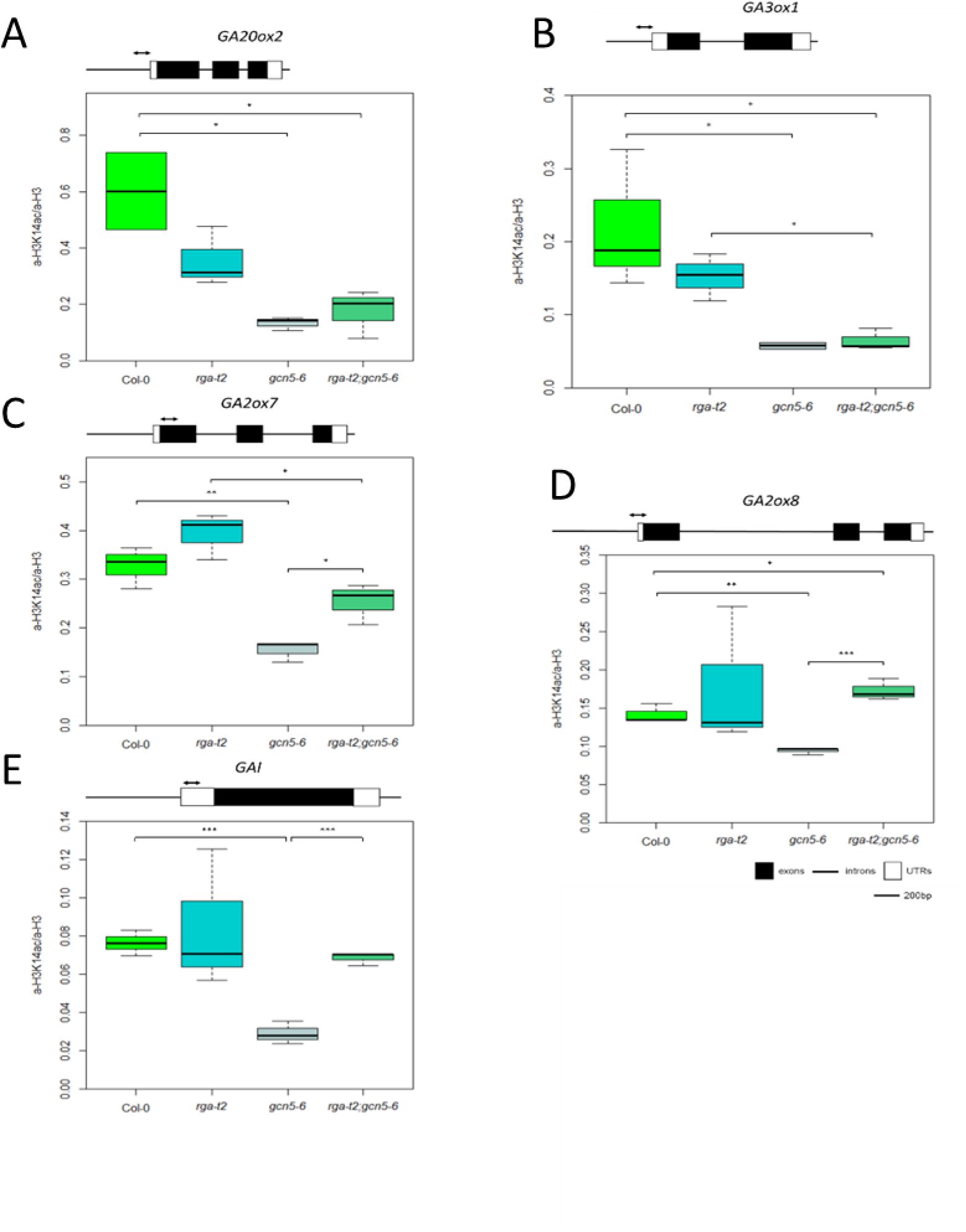
The acetylation status of histone 3 lysine 14 (H3K14) in the gibberellin biosyntheses genes *GA20ox2 (A)* and *GA3ox1 (B)*, gibberellin catabolism genes *GA2ox7 (C)* and *GA2ox8 (D)*, and DELLA gene *GAI (E)*. Col–0, *rga–t2*, *gcn5–6*, and *rga–t2;gcn5–6* inflorescences were harvested for chromatin immunoprecipitation using antibodies against acetylated H3K14 and H3. Error bars show the standard deviation. Asterisks show statistical significance compared to Col–0 and *rga–t2;gcn5–6* double mutant, using Student’s *t*-test: *, *P* < 0.05, **, *P* < 0.01, and ***, *P* < 0.001.

### Mutation in *RGA* did not suppress *ada2b* phenotypes

In Arabidopsis, non-functional *ada2b* mutants exhibit developmental problems such as dwarfism, delayed root growth, flowers with short petals and stamens, increased infertility (Vlachonasios et al. 2003) and reduced expression of GA biosynthesis genes (Fig. 2B). Many of these phenotypes are similar to *gcn5* mutants since ADA2b is physically associated with GCN5 (Mao et al. 2006) and could be related to plant responses to GA, suggesting potential problems in GA signaling. Therefore, we characterise *ada2b-1; rga-t2* and *ada2b-1;rgl2-1* double and *ada2b-1; rga-t2;rgl2-1* triple mutants to explore if RGA alone or with RGL2 could partially suppress the *ada2b* phenotype. The supplemental Fig. S6A clearly shows that the triple *ada2b-1;rga-t2;rgl2-1* and the double mutants *ada2b-1;rga-t2* and *adab-1;rgl2-1* display a phenotype similar to that of *ada2b-1*, characterized by short root length compared to the wild-type seedlings. In addition, the mutants have an increased number of secondary roots and elongated hypocotyl. Later in the plant development, during the flowering period, both the double mutants *ada2b-1;rga-t2*, and *ada2b-1;rgl2-1* and the triple mutant *ada2b-1;rga-t2;rgl2-1* display a dwarf phenotype similar to *ada2b-1* plants (Supplemental Fig. S6B). The inflorescence elongation was severely limited relative to the wild-type plants. These results indicate that the absence of *RGA* and *RGL2* does not reverse the *ada2b-1* phenotype. Therefore, ADA2b function is not suppressed by RGA, suggesting a distinct regulation of ADA2b and GCN5 in the GA signaling pathway.

## Discussion

This manuscript explored the role of histone acetyltransferase GCN5 and the transcriptional adaptor ADA2b in gibberellin responses. We found that GCN5 regulate the expression of genes involved in GA biosynthesis, catabolism and signaling in Arabidopsis by modulating the histone acetylation. Furthermore, we showed that the DELLA protein RGA partially suppresses the GCN5 function in flower development and stamen elongation independent from ADA2b.

Histone acetyltransferase GCN5 and the associated coactivator ADA2b are involved in diverse developmental processes, including root growth, cell elongation, trichome differentiation, floral initiation, apical meristem function and floral reproduction (Vlachonasios et al. 2021). Many of those processes could arise from defects by GA signaling. Gibberellins affect many biological processes, including seed germination, cell elongation, transition to flowering and flower development (Sun 2008). Our data suggest that GCN5 affect histone acetylation (H3K14Ac) in the promoter of genes involved in the last step of GA biosynthesis and members of GA2 oxidases involved in GA catabolism. These effects are correlated with the gene expression, suggesting that GCN5 is a positive regulator of GA homeostasis in the early-formed flowers. The effect of GCN5 on GA-inactivating genes is tissue and gene-specific since *GA2ox4* and *GA2ox8* are upregulated in *gcn5* mutants, whereas *GA2ox6* expression was downregulated in hypocotyls. *GA2ox6* was suggested as a target of the PAGA complex, which contains GCN5 and ADA2a in 12-day-old seedlings (Wu et al. 2023).

Without gibberellins, the DELLA proteins positively regulate the late GA-biosynthesis genes and the receptors family GID1, whereas they negatively regulate the GA-inactivating genes (Daviere and Achard 2013). Therefore, we explored genetically the effect of RGA on GCN5 function by characterising the double mutants. Our data suggest that the RGA could partially suppress many *gcn5* phenotypes during the life cycle of Arabidopsis. In the seedlings, RGA suppresses the positive role of GCN5 on hypocotyl elongation, which is correlated with the effect on *GA3ox1*, *GA2ox4*, and *RGL2* gene expression. During reproductive stages, RGA suppresses the GCN5 effect on the primary inflorescence growth, especially in the late developmental stages and the formation of secondary inflorescences, restoring the bushy appearance of *gcn5* mutants and suggesting that RGA mediates the positive role of GCN5 on apical dominance. Indeed, RGA represses GA-induced apical dominance (Dill and Sun 2001).

GCN5 is essential in flower development, especially in reproductive organs, stamens and gynoecium (Cohen et al. 2009; Poulios and Vlachonasios 2018; Vlachonasios et al. 2021). GCN5 promote stamen filament elongation in the early-formed flowers (Cohen et al. 2009; this study) by affecting the expression and the histone acetylation levels on late-stage biosynthetic genes, *GA20ox2* and *GA3ox1*, as well as the GA-inactivating genes. *GA3ox1* is expressed in the early flowers’ stamen filament (Hu et al. 2008). Our data suggest that the positive role of GCN5 on stamen filament growth is suppressed by RGA action. This scenario does not arise from changes in gene expression and histone acetylation levels of GA biosynthesis. However, the effect of RGA is concentrated on the expression level of GA catabolism genes, GA receptor GID1b and the DELLA protein GAI, suggesting that the GCN5 effect on those genes is mediated by RGA action.

Moreover, the effect of GCN5 on histone H3K14 acetylation levels in the *GAI* promoter is restored by RGA either by recruiting another histone acetyltransferase or by inhibiting the action of a histone deacetylase. During flower initiation, the H3K14 acetylation levels in the GAI locus are positively affected by ADA3a, a component of the SAGA complex (Poulios et al. 2021). HDA15 is recruited in the promoter GA biosynthesis genes and repressed their expression (Zheng et al. 2022). Beyond the gibberellin effect on stamen filament elongation, another plant hormone, jasmonic acid, is also involved (Marciniak and Przedniczek 2019). RGA and RGL2 are critical for the inhibition of stamen development (Cheng et al. 2009) by interacting with transcription factors MYB21 and MYB24, acting as a central synergistic node of GA and JA signaling on stamen filament interaction (Huang et al. 2020). The opposite activities of GCN5 and HDA6 regulate TPL acetylation and repressor activity to determine the transcription of JA-responsive genes (An et al. 2022). This scenario could also explain the effect of RGA on GCN5 action on stamen elongation.

Our data also suggest that RGA does not repress several GCN5 actions. Those include the promotion of root elongation in seedlings, the effect of GCN5 leaf serration patterning and development, and the positive role of fruit elongation and development. Interestingly, RGA and RGL2 could not suppress the ADA2b action, suggesting that ADA2b act downstream of DELLA action. Furthermore, GCN5 and ADA2b could have distinct mechanistic regulations in GA signaling.

Although it is unclear if there is a physical interaction between RGA and GCN5 or other members of the SAGA complex, recently RGA was found to interact with H2A to form a complex between transcription factors, RGA and H2A (Huang et al. 2023). In the future, it is necessary to provide mechanistic evidence of the possible DELLA-Histone acetylation interaction and how this regulates developmental stages, tissues and cell type differentiation.

In conclusion, our findings add essential evidence to our current understanding of mechanisms involving the interaction of gibberellins and histone acetylation. We have shown that the GA signaling repressor RGA suppresses the histone acetyltransferase GCN5 action on stamen filament elongation by affecting histone acetylation-mediated gene expression on GA catabolism and signaling (Fig. 9A and B). Additional molecular and genetic studies may further dissect the role of GCN5 in the development of reproductive organs, and their complex interaction and biochemical analyses may reveal mechanisms by which histone acetylation affects GA signaling.

**Figure 9.**
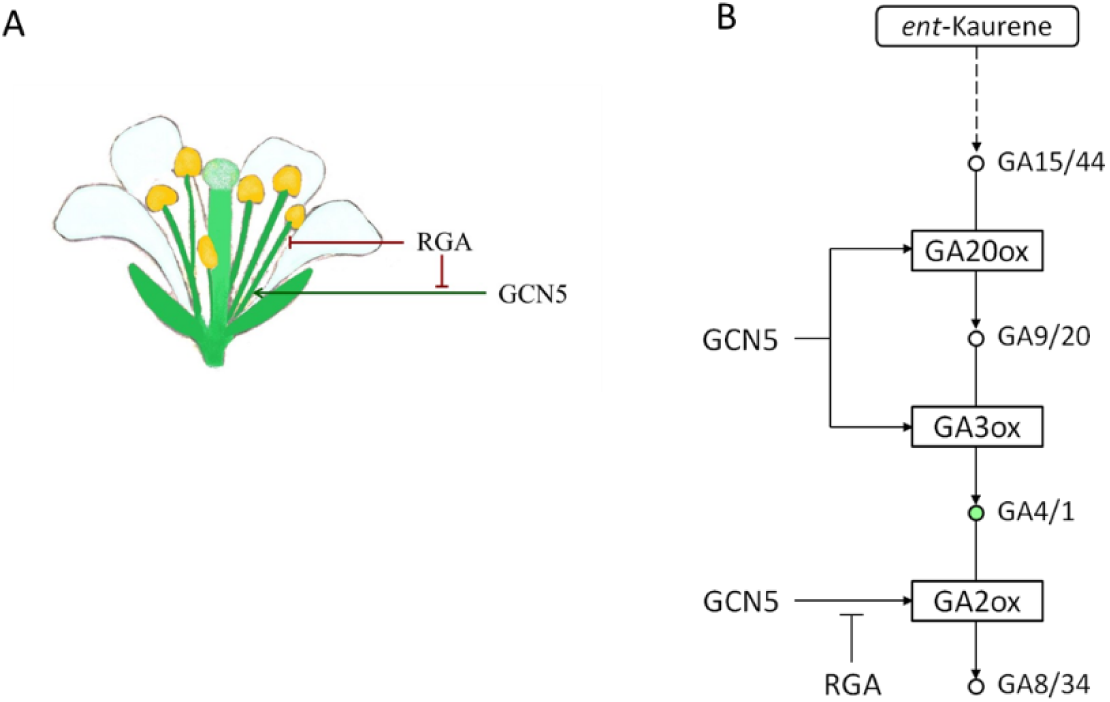
The genetic interaction between *RGA* and *GCN5* on stamen filament growth (A). GCN5 is a positive regulator of GA biosynthesis and catabolism. RGA suppresses GCN5 action on GA2 oxidases (B).

## Materials and Methods

### Plant material and plant growth conditions

In this study, *Arabidopsis thaliana* ecotypes Col-0 and Ws-2, as well as single mutants *ada2a-3, ada2b-1, gcn5-1, gcn5-6/hag1-6* and *prz1-1* (Sieberer et al. 2003; Vlachonasios et al., 2003; Long et al., 2006; Hark et al, 2009), were used. Seeds were first disinfected with 30% bleach for 5 min and then stratified at 4°C for three days. The seeds were sown in Petri dishes containing Gambrog B5 nutrient medium (Douchefa, Haarlem, Netherlands) with pH 5.7, 1% sucrose and 0.8% w/v phyto-agar (Douchefa, Haarlem, Netherlands) and then transferred to a growth chamber with a constant temperature of 20°C and a constant light intensity (100μmol*m^-2^*s^-1^). The plant growth photoperiod was 16 hours of light and 8 hours of darkness.

### Hypocotyl and root elongation measurements

After five days of germination, seedlings were transplanted into Petri dishes containing Gamborg B5 medium (Duchefa, Haarlem, Netherlands) and different concentrations of gibberellic acid (0 μM GA_3_, 2.5 μM GA_3_ and 10μM GA_3_) without sucrose. These petri dishes were placed perpendicular to the light. Photos were taken for four consecutive days and processed with the ImageJ (NIH, Maryland, USA). The experiments were repeated thrice, and 20 seedlings per genotype per treatment were used.

### Genetic analysis

The *Arabidopsis thaliana* (L) Heynh. *gcn5-6* (*hag1-6*) and *ada2b-1* mutants were previously described in (Vlachonasios et al., 2003, Long et al., 2006). The *rga-t2* was obtained by the Nottingham Arabidopsis Stock Centre (NASC) and described (Lee et al., 2002). The *gcn5-6;rga-t2* and the *ada2b-1;rga-t2* double mutants were created using pollen from *rga-t2* homozygous mutants to fertilise *gcn5-6* and *ada2b-1* gynoecium. The resulting F1 generation was self-fertilised, and the segregating F2 population was genotyped using PCR-based methods. The enzyme ExTaq DNA polymerase (Takara, Japan) was used with the primers listed in Supplemental Table S4 to confirm the double mutants. The double mutants were backcrossed to Ws-2 or Col-0 background for at least four generations. Kanamycin resistance of *ada2b-1* alleles was used when applicable to facilitate selection.

### Gene expression assays

For RT-qPCR expression analysis, whole inflorescences from Col-0, *rga-t2, gcn5-6* and *rga-t2;gcn5-6* plants were collected when the first one or two open flowers emerged and flash-frozen in liquid nitrogen. The frozen tissue was preserved at −70^ο^ C. Five different harvests were made, and three were used for RNA extraction using the Nucleospin^®^ RNA Plant kit (Macherey-Nagel, Duren, Germany). RNA quality and quantity were assessed by 1.5% gel electrophoresis and NanoDrop 2000 (Thermo Fischer Scientific, Waltham, MA, USA). The PrimeScriptTM 1st strand cDNA synthesis kit (Takara, Shiga, Japan) was used for reverse transcription. In three independent biological repeats, reverse transcription was performed using 0.5 μg of total RNA. Quantitative reverse-transcription polymerase chain reactions (RT-qPCRs) were prepared with the AMPLIFYME SG Universal Mix (AM02) (BLIRT SA, Gdansk, Poland) or the Luna® Universal qPCR Master Mix (New England Biolabs, Ipswich, MA, USA) at the ABI StepOnePlus™ system (Applied Biosystems, Foster City, CA, USA). Three technical repeats were run for each sample. The *At4g26410* or the *PDF2* genes were used as endogenous controls (Supplemental Table S4). Data were analysed with the ΔΔCt method using StepOne Software 2.1. Statistical analysis was performed using R.

### Chromatin Immunoprecipitation assays

Whole inflorescences of Col-0, *rga-t2*, *gcn5-6* and *rga-t2;gcn5-6* plants were collected when the first one or two open flowers emerged. The tissue was fixed in 1% formaldehyde under vacuum for 15 minutes, and the crosslinking reaction was terminated with 0.125M glycine for 5 minutes under vacuum. Samples were stored at −70^ο^ C. Approximately 300mg of tissue from each genotype were ground in liquid nitrogen to a fine powder, and nuclei were extracted and lysed in the presence of 1% SDS. Chromatin was sheared into 200 to 1000bp fragments using Fisherbrand™ Model 505 Sonic Dismembrator (Fischer Scientific, Waltham, MA, USA) with the following parameters: sonication for 10s and stopping for 50 s at 50% power five times. Chromatin was diluted ten times before the immunoprecipitation, with antibodies against acetylated histone H3K14 (Anti-Histone H3 (Lys14), EMD Millipore #07-353, Burlington, MA, USA) and non-acetylated histone H3 #ab1791 (Abcam, Cambridge, UK). The precipitation was performed using agarose-protein A beads (Cell Signaling, Danvers, MA, USA). The elution of chromatin attached to the beads was made at 65 °C with 1% SDS and 0.1M NaHCO_3_. Formaldehyde crosslinking was reversed in the presence of 200mM NaCl at 65 °C overnight, followed by proteinase K (Sigma-Aldrich, St Louis, MI, USA) treatment. The DNA was isolated using a commercially available PCR clean-up kit (Macherey-Nagel, Duren, Germany). Immunoprecipitated DNA was diluted in water and analysed with qPCR using specific primers (Supplemental Table S4). Luna® Universal qPCR Master Mix (New England Biolabs, Ipswich, MA, USA) and the ABI StepOnePlus™ system (Applied Biosystems, Foster City, CA, USA) were utilised for the qPCR assays. Ten-fold serial dilutions of input Col-0 were used to create a standard curve. All data obtained by q-PCR were presented as a percentage of input. The value of each immunoprecipitated sample was normalised to the input. The ratio of acetylated H3K14 to H3 values of each genotype is presented in the graphs. The immunoprecipitation assays were performed in three independent biological repeats. Statistical significance was calculated using Student’s t-test in R.

## Supplementary Data

Supplementary Figure 1. The effect of GA_3_ on root elongation of Ws-2, *ada2b-1*, *gcn5-1*, Col-0, *prz1-1*, *gcn5-6*, and *ada2a-3* seedlings.

Supplementary Figure 2. Col-0, *rga-t2*, *gcn5-6* and *rga-t2;gcn5-6* mutant plants after 57 days of growth.

Supplementary Figure 3. A) *gcn5-6* and *rga-t2;gcn5-6* mutant plants after 57 days of growth and B) *gcn5-6* mutant plants after 90 days of growth.

Supplementary Figure 4. *rga* suppress *gcn5* infertility.

Supplementary Figure S5. Expression of gibberellin biosynthesis gene *GA3ox1* (a), gibberellin catabolism genes *GA2ox4*, *GA2ox6*, and *GA2ox8* (b,c and d) GA-receptors *GID1a* and *GID1b* (e and f), and DELLA genes *GAI* and *RGL2* (g and h) in hypocotyl of *Arabidopsis thaliana*.

Supplementary Figure S6. Characterization of *ada2b;rga*, *ada2b;rgl2*, double mutants and *ada2b;rga;rgl2* triple mutants at seedlings stage, and after 42 days of growth in long days conditions.

Supplementary Table S1. The effect of *GCN5* and *RGA* on the number of lateral inflorescences.

Supplementary Table S2. The effect of *GCN5* and *RGA* on the number of secondary inflorescences.

Supplementary Table S3. The effect of *GCN5* and *RGA* on the number of stamens in early flowers.

Supplementary Table S4. Primers used in this study.

## Acknowledgements

We would like to thank Professor Amy Hark (Muhlenberg College, Allentown, USA) for stimulating discussion and critical reading of the manuscript.

## Funding

This work was partially funded by the Hellenic Foundation for Research and Innovation Research project MIS 3026 to KV. Erasmus^+^ scholarship was granted to DT and ZS. The types of equipment used in this study were obtained by the project “Upgrading the plant capital” (MIS 5002803), which is implemented under the Action “Reinforcement of the Research and Innovation Infrastructure”, funded by the Operational Programme “Competitiveness, Entrepreneurship and Innovation” (NSRF 2014–2020)and co-financed by Greece and the European Union (European Regional Development Fund).

### Author contribution

Designed the research (KV); performed research (CB, SP, DT, ZS, MCK, AK); analyzed data (CB, SP, KV); or wrote and edited the mauscript (KV, JD).

